# SK3-overexpression drives sex-dimorphic working and social memory impairment in mice

**DOI:** 10.1101/2020.07.31.230656

**Authors:** Alline C Campos, Franciele F Scarante, Sabine Martin, Marcio Lazzarini, Doris Hermes, Maria A De Souza-Silva, Luis A Pardo, Walter Stühmer, Elaine A Del-Bel

**Affiliations:** Department of Pharmacology, Ribeirão Preto School of Medicine, University of São Paulo, Ribeirão Preto, Brazil; Department of Molecular Biology of Neuronal Signals, Max Planck Institute of Experimental Medicine, Göttingen, Germany; Center Nanoscale Microscopy and Molecular Physiology of the Brain (CNMPB), Göttingen, Germany; Institute of Experimental Psychology-University of Düsseldorf, Düsseldorf, Germany; Oncophysiology Group, Max Planck Institute of Experimental Medicine, Göttingen, Germany; Department of Basic and Oral Biolgy, Dental School of Ribeirão Preto, University of São Paulo, Ribeirão Preto, Brazil

**Keywords:** memory, sex differences, SK3 channels, neuroplasticity, adult hippocampal neurogenesis, estrogen signaling

## Abstract

Although sex differences in memory tasks dependent on hippocampal function have been described in several species, including rodents and humans, the exact mechanisms involved remain debatable. The function of the small-conductance Ca2+-activated K+ channel type 3 has been associated with cognitive deficits, and its overexpression in male mice (T/T) induces shrinkage of the hippocampus. Here we describe that opposite to the observation in males, in female mice, SK3-induced-reduction in the volume of the hippocampal formation does not interfere with working and social memory performance. Male, but not female T/T mice showed decreased adult hippocampal neurogenesis and down-regulation of the expression of the genes related to Akt/mTOR and MAP kinase pathways. T/T male mice exhibit impaired estrogen and Neurogulin 1 signaling. An increased number of filopodia spines is observed in the dentate gyrus (DG). Our results suggest a fine-tune modulation of SK3 expression participates in the sex-dependent function of the hippocampus via estrogen signaling and neuroplasticity in the DG. Our results reinforce the importance of testing male and female mice while conducting experiments with transgenic mice.

## 1.0 Introduction

In the field of biomedical science, studies using genetically modified rodents have generated essential data that continuously help our society to understand the nature of diseases and the discovery of new treatments. A representative part of these studies was conducted only on male mice, ignoring possible sex differences, and biased the interpretation of traits and disease phenotypes (Karp et al., 2017). This panorama is not different in Neuroscience.

The function of brain structures of all animals, including humans, are developed and refined under the influence of sex and the environment (Levine 1966, Cahill, 2006). Sex dimorphism impacts the expression of emotion, memory, vision and hearing functions, pain perception, spatial navigation, and the action and the levels of neurotransmitters and hormones during health or disease (Cahill, 2006).

One of the central brain structures that are sexually dimorphic is the hippocampus (Cahill, 2006; Mizuno and Giese, 2010; Koss and Frick, 2017). This brain region is extensively associated with learning processes, memory, and emotions, presenting remarkable anatomical and neurochemical differences between males and females (Madeira & Lieberman, 1995). However, the mechanisms involved in how sexual dimorphism contributes to the memory processes dependent on hippocampal function are far from clear. Memory is a complex process influenced physiologically (aging, neurotransmitters, intracellular pathways) and exogenously (stress, environment) (Koss & Frick, 2017).

Small-conductance calcium-activated potassium channels (SK) control calcium homeostasis and modulate membrane excitability (Sah, 1996). They contribute to adjusting the duration and amplitude of the action potential afterhyperpolarization (AHP), critical events for the correct function of neuronal somatic excitability (Pedarzani et al., 2005). These potassium channels also play an essential role in the regulation of neuron firing patterns, neurotransmitter release, and synaptic plasticity, given their close relationship with changes in calcium concentrations (Pedarzani et al., 2005).

The type 3 SK channel (SK3 or Kca2.3) is encoded in humans by the KCNN3 gene and maps to chromosome 1q21, a region linked to schizophrenia. Blank and colleagues (2003) demonstrated that SK3 is more expressed in the aged mouse brain, contributing to the decline of cognition and long-term potentiation (LTP). Grube and colleagues (2011) suggested that male mice’s overexpressing SK3 channels exhibited deficits in spatial learning and the expression of fear conditioning. More recently, our group has shown that transgenic SK3-T/T male mice exhibit working memory deficits, impaired LTP, and significantly shrinkage of the hippocampal formation (Martin et al., 2015).

SK3 channel is sensitive to small concentrations of Ca2+. Changes in the cytosolic free Ca2+ concentration play a central role in many fundamental cellular processes like neurotransmitter release, cell proliferation, differentiation, gene transcription (Berridge et al., 2003), and cell signaling pathways, such as Akt/mTOR and MAPK (Hoyer-Hansen et al., 2007). All the described processes are related to memory formation within the hippocampus.

Interestingly, the mRNA expression of SK3 channels seems highly regulated by estrogen (Pierce & England, 2010). Estrogen is one of the sex hormones believed to participate and contribute to dimorphism observed during memory formation, including the hippocampal-dependent process (Mizuno & Giese, 2010). In the hippocampus of mice, SK3 channels are expressed in both pre- and postsynaptic neurons (Tacconi et al., 2001; Ballesteros-Merino et al., 2014) in the CA3 and the dentate gyrus (DG-Sailer et al., 2004).

The DG, mainly its granular cell layer, is a highly dimorphic structure during adolescence and adulthood. Under the influence of testosterone, which is converted into estradiol in the brain, granule neurons change their size and volume during spatial memory task performances (Roof, 1993). Within the hippocampus, the DG is the main entrance gate for information and its vast capacity of neuroplasticity (such as LTP, dendritic remodeling and adult hippocampal neurogenesis) confers to this area relevant roles in the formation of new memories, and in encoding and retrieval of contextual clues in situations where associative learning is needed (Bernier et al., 2017).

However, the relationship between the SK3 function, DG plasticity, and dimorphism during the memory process has not been addressed. Therefore, in the present study, we hypothesized that overexpression of SK3 channels contributes to the mechanism of sex dimorphism associated with cognitive processes mediated by the DG within the hippocampus. Our results demonstrate that despite the hippocampal shrinkage, female, but not male, transgenic mice overexpressing SK3 channels can perform memory tasks and retain intact neuroplasticity in the DG (adult hippocampal neurogenesis), intact V-akt murine thymoma viral oncogene (Akt)/ mechanistic target of rapamycin (mTOR) and mitogen-activated protein kinase (MAPK) pathways functions. In our working hypothesis, we believe that estrogen signaling (impaired in male SK3 mice, but intact in female) is counterbalancing and maybe controlling the SK3 function to maintain memory function in the hippocampus of female mice.

## 2.0 Results/Discussion

### 2.1 Female, but not male SK3-T/T mice, can perform short- and long-term memory tasks despite hippocampus shrinkage

Previous work of our group has demonstrated that male T/T mice exhibited: deficits in spatial memory acquisition in the Morris Water Maze; impaired short-term memory tested in Novel Object Recognition (NOR) test, and less expression of freezing behavior in a fear conditioning paradigm (Grube et al., 2011; Martin et al., 2015).

Here we showed that despite SK3 transgenic mice of both sexes exhibit hippocampal shrinkage (Supplementary Figure 1), only male T/T mice presented impaired short-(t(17)= 8.31, p<0.001) and long-term memory (t(17)= 8.06, p<0.001) in the NOR test (Figure 1A-D). These deficits are evidenced by the significant reduction in the percentage of time exploring the novel object 1.5 and 24 hours after the training session in the NOR test.

**Figure 1.**
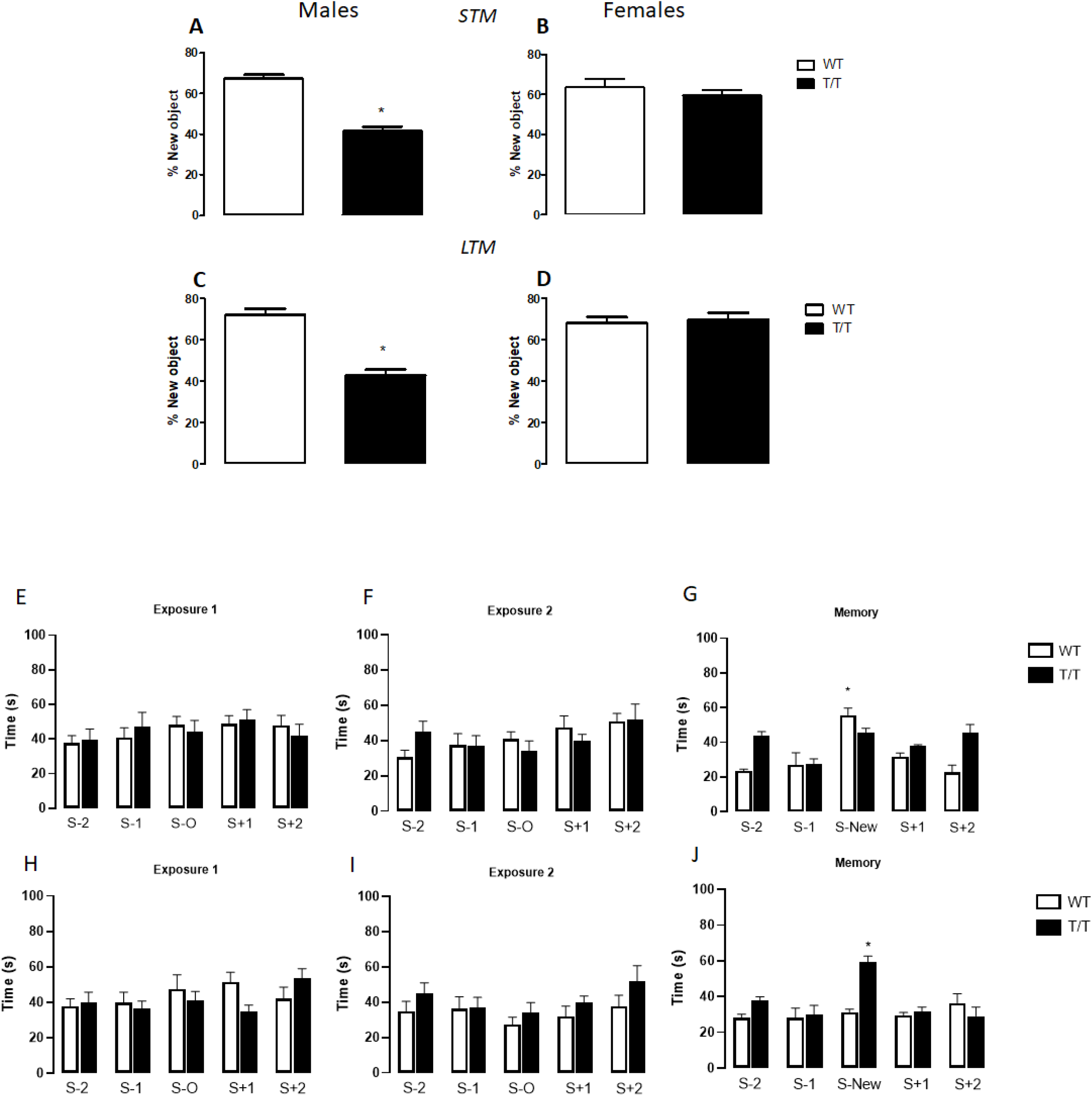
SK3 overexpression leads to sex-dimorphic working and social memory impairment. A and C- short- and long-term memory tasks performed by male mice in the Novel Objects Recognition Test (NOR); B and D- short- and long-term memory tasks performed by female mice in the NOR (n=10 per group). E-J: results derived from the social box test. E, F, H, and I: pre-test/habituation sessions of male and female, respectively. G and J: Memory retention test of male and female, respectively. (N= 10 per group). Bars represent mean+/− standard error mean (SEM). * represents differences to the WT group whereas # represents differences between trials of the same genotype; p< 0.05.

In addition to the NOR, we also tested if SK3 overexpression-induced hippocampal shrinkage would decrease social memory in female and male T/T mice. Similar to the behavioral alterations found in NOR, another memory paradigm, in the memory phase of the SocioBox (Krueger-Burg et al., 2016) test (Figure 1E-1J) male T/T mice fail to perform the memory task. In contrast, the female T/T spent significantly (p = 0.013) more time with the stranger mouse (S new) compared to the other stimulus mice (S-2, S-1, S+1, S+2). No effects of genotype or sex differences were detected on the preference for any position during exposure 1 (figure 1E and 1H) or exposure 2 (figure 1F and 1E).

The inherent capacity that mammals retain to distinguish between different individuals and maintain the distinction over time has a critical role in memory itself, but also on social cognition perspectives and skills. The hippocampus is one of the most critical regions responsible for the disambiguation of overlapping sequences and similar stimuli (Agster et al., 2002; Ross et al., 2013). During the NOR and the SocioBox test, mice use that hippocampal ability to discriminate a familiar object or housemate, from a new one. Even with a reduced volume of the hippocampal, female SK3 mice are somehow able to perform working memory and discriminate against social cues.

Interestingly, the dimorphic sex effects of SK3 overexpression seem to be specific for memory processes, as indicated by our results in other animal models not causally related to the mnemonic process. For instance, in the prepulse inhibition test (PPI-Supplementary figure 2), a model frequently used to detect sensorimotor gating impairment (Swerdlow and Geyer, 1998) that also involves hippocampal function, we did not observe any behavioral changes in SK3 T/T mice (both sexes). (Supplementary figure 2). The PPI paradigm has been used to test the involvement of the hippocampus, genes, and candidate drugs in the sensory gating deficiencies observed in schizophrenic patients (Kumari, 2011, Le Pen and Mureau, 2002). In a previous study of our group, CAG repeats in the code-region of KCNN3was associated with reduced function of SK3 channels and correlated with better cognitive performance of schizophrenic patients (Grube et al., 2011). A meta-analysis study published in 2003 has concluded that CAG region repetition was not correlated with a high risk of schizophrenia (Glatt et al., 2003).

**Figure 2.**
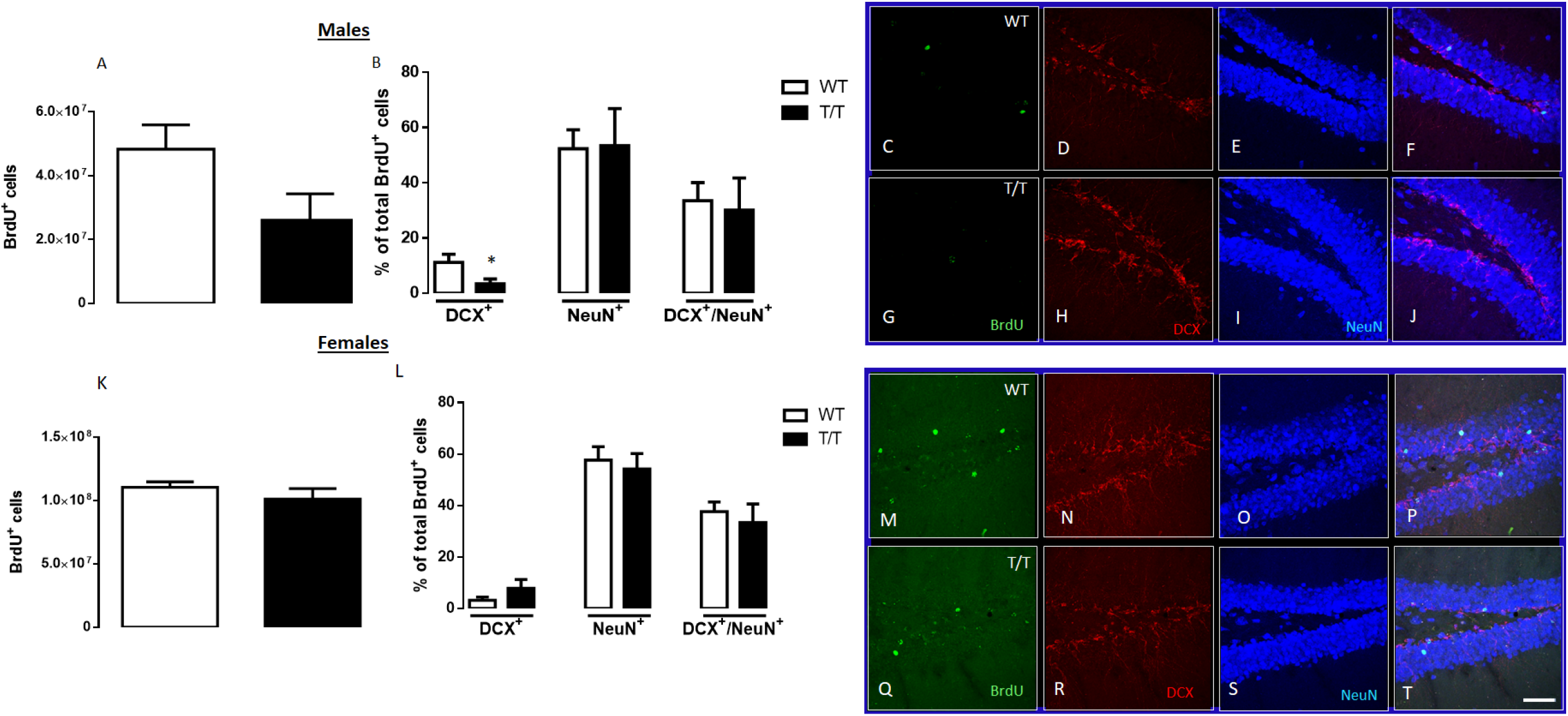
Male T/T mice, but not females, exhibit decreased hippocampal neurogenesis. A and K-BrdU positive cells in female and male mice, respectively. BrdU positive cells were analyzed 21 days after BrdU injections. B and L: double and triple labeling of BrdU+ Doublecortin (DCX), BrdU+ NeN, or BrdU+ DCX +NeN. C, G, M, and Q- Representative images of BrdU, D, H, N, and R, Representative images of DCX. E, I, O, and S: Representative images of NeuN. F, J, P, and T: representative images of the triple labeling. In the graphics, bars represent mean+/− SEM. In the illustrative images, the bar represents 50um (pictures were taken at a magnification of 20X). * represents the difference relative to the WT group. N= 5/group.

Also, given the relationship between the hippocampus and major depressive disorder, we evaluated if the reduced volume of the hippocampus observed in T/T mice would affect depressive-like behaviors. Jacobsen and colleagues (2008) suggested that the genetic ablation of SK3 induces a reduction in the time spent immobile in the tail suspension test (TST) and the forced swimming test in a sex-independent manner. In our study however, we demonstrated that SK3 overexpression also induces an antidepressant-like phenotype in the TST (Supplementary figure 2, 1F and 1G; males-t(18)= 4.98, p<0.001; female-t(18)= 3.30, p<0.01). T/T mice exhibit an increased preference for sucrose when compared to food-deprived WT animals (males-t(18)= 2.38, p<0.05; female-t(18)= 4.09, p=0.001) (Supplementary figure 2). T/T animals exhibit no change in the level of corticosterone as compared to wild type animals (Supplementary Figure 3). Our results also demonstrate that, besides decreased passive coping behavior, SK3 could also be involved in anhedonia-like behaviors. These results may implicate SK3 in the mechanisms related to mood disorders, such as Major Depressive Disorders.

**Figure 3.**
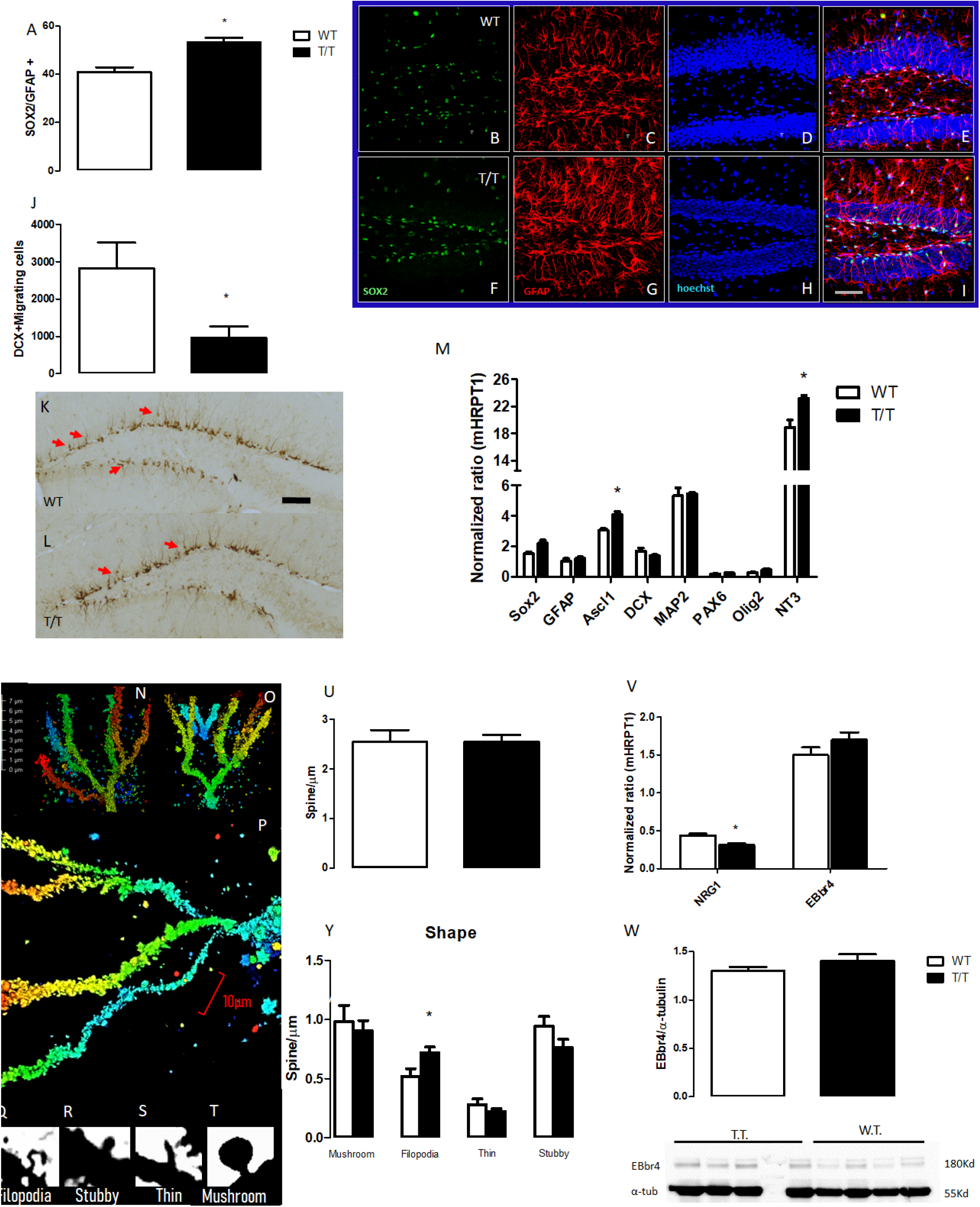
SK3-overexpression induces an increased number of Type 1 progenitors (SOX2+ GFAP+) and filopodia spines, impaired cell migration, and Neuregulin-1 signaling in MALE mice. A-I: results from the double-labeling analysis of cells expressing SOX2+GFAP in the subgranular zone of the dentate gyrus (DG). J, K, and L: display the results related to doublecortin positive cells migrating towards the granular zone of the DG. Arrows indicate cells in the process of migration (K and L). M-results from the qPCR analysis of the DG of the hippocampus (N=3/group). N-X: results and representative images of the analysis of the number and shape of dendritic spines in the granular neurons of the hippocampus. V and Y: qPCR and WB for Neuregulin (NGR1) and its receptor EBbr4. Bars represent mean+/− SEM. * indicates a difference with respect to the WT group. N=5 per group except for the qPCR analysis).

Despite the complex behavioral alterations and the sexual dimorphism described in the memory performance of T/T mice, we interpret that the imbalance promoted by overexpression of SK3 channels is affecting the synaptic function within the hippocampus. In the long term, this would culminate in the drastic reduction of the hippocampal volume observed in these transgenic mice. Ion channels are dynamic modulators of all aspects of the cell biology of mature, immature, and neural precursor cells (NPC). Neuronal adaptation might be mediated by slow afterhyperpolarization (sAHP), generated by potassium currents developed after spike firing (Madison and Nicoll 1984; Saar et al. 2001).

It has been shown that decreased post-burst AHP correlates with a worse performance during learning and memory tasks (Disterhoft et al. 1996; Saar et al. 1999). SK channels limit burst frequency because, during a train of action potentials, the duration of the sAHP period is increased, rendering the cells unable to reach another action potential threshold (Sah, 1996). Overexpression of SK3 would make the threshold for SK activation more sensitive to Ca2+ currents, because smaller Ca2+ concentrations would activate a more significant number of SK3 channels to limit burst frequency, increasing the time of sAHP and therefore delaying neuronal firing (Deignan et al., 2012).

Several studies support the idea of decreased neuronal firing during aging. In the senescent hippocampal neurons, the magnitude of the Ca2+-dependent, K+-mediated AHP is facilitated (Kumar and Foster, 2007). The amplitude of the AHP is consistently increased and prolonged in the hippocampal CA1 pyramidal neurons during aging (Landfield & Pitler, 1984; Kumar and Foster, 2007). Blank et al. (2003) demonstrated increased expression of the SK3 channel in the aged hippocampus, which contributes to LTP deficits and age-dependent decline of learning and memory. Accordingly, SK3 overexpression would accelerate hippocampal aging, which may increase hippocampal cell death via apoptosis (increased Caspase 3 expression in CA1-Supplementary figure 3) and inducing the memory impairment observed in male T/T (Grube et al., 2011, Martin et al. 2017). However, those arguments still do not explain why female T/T can perform memory tasks despite their shrunk hippocampus.

### 2.2 SK3 overexpression decreases adult hippocampal neurogenesis in male, but not female, mice

To further investigate the mechanism that could explain why SK3 overexpression does not affect memory performance in female mice, we analyzed if SK3 would affect hippocampal neuroplasticity equally in males and females. The hippocampus is one of the regions that retain the capacity to produce new neurons in the adult brain. To characterize the role of SK3 channels on the process of adult hippocampal neurogenesis in both female and male mice, we performed confocal microscopy analysis for the quantification of 5-Bromo-2’-deoxyuridine (BrdU)-positive cells, co-labeled with Doublecortin (DCX), a marker for migrating cells and immature neurons, and NeuN, a marker for mature neurons.

Our results suggest that SK3 overexpression changes the dynamics of the formation of new neurons in the DG of the hippocampus only in males. Surprisingly, despite the hippocampal shrinkage, female T/T show intact hippocampal neurogenesis. Recently, Berdugo-Vega and colleagues (2020) have suggested that adult neurogenesis can improve hippocampal activity, “rejuvenating” strategies of learning and memory. Also, it was shown that in senescent DG, facilitation of adult neurogenesis might rescue hippocampal function (Kempermann et al., 1998). Accordingly, adult-born neurons seem to spike, engage, and connect faster in the DG, which contributes to better execution of memory processes (Snyder et al., 2009). This process seems to be conserved in a female, but not male, T/T mice. Conversely, male T/T mice presented 85% less BrdU-positive cells (corrected by the dentate gyrus size, see methods) than their WT littermates (t(8)= 2.38, p=0.051 Figure 3 A and C), indicating that SK3 channels might play an important role during the survival phase of adult neurogenesis. The percentage of BrdU-positive cells that also expressed DCX was 227% larger in the WT mice (t(8)= 2.45, p<0.05, figure 2 D and I), indicating that SK3 overexpression leads to a lower number of newly-generated neurons (Type 2b and type 3 cells) during differentiation (t(8)= 2.38, p<0.05). No statistical differences were found in the analysis of co-labeled cells BrdU+/DCX+/NeuN+ or BrdU+/NeuN+ (Figure 2, F and G, J, and K). This apparent contradiction might reflect the time-window of our analysis (21 days instead of 28 or more-Kempermann et al., 1998) and the degrees of maturation of new neurons (Aimone et al., 2010). In the case of female T/T mice, no changes in the number of BrdU+, BrdU+DCX+, BrdU+/DCX+/NeuN+, or BrdU+/NeuN+ positive cells were observed (Figure 3, N-U).

Limited information is available about the role of SK3 in neural stem cells and adult hippocampal neurogenesis. SK3 is the most abundant KCa channel in precursor cells, predominantly found in nestin-positive cells (Liebau et al., 2007). In rat mesenchymal cells, intermediate conductance KCa channels reduce cell proliferation by accumulating cells at the G0/G1 phase (Tao et al., 2007). Our results support the idea that the SK3 channel controls DCX (a microtube-associated protein) expression, decreasing cell migration and forming type 2b and type 3 cells in the adult hippocampus. SK3 channels co-localize with nWASP (neural Wiskott-Aldrich syndrome protein), a member of the filopodial initiation complex (Carlier et al. 1999; Yoo et al., 2006). Liebau and coworkers (2011) have demonstrated in vitro that SK3 channel/nWASP/Abi-1 complex controls prolongation and neuronal differentiation during embryonic stages.

Our results demonstrated that T/T mice (males) present an increased number of double-labeled Glial Fibrillary Acidic Protein (GFAP) and (sex-determining region Y)-box 2 (SOX2)-positive cells (t(8)= 4.52, p<0.01, Figure 3A and figure 3-C and G, representative images) in the subgranular zone of the hippocampus, indicating a possible involvement of SK3 retaining type 1 precursors phenotype – possibly quiescent cells (Figure 3E and 3I). Also, our analysis revealed that SK3 male mice have fewer DCX+ cells migrating from the subgranular to the granular zone of the DG (Figure 3J, and 3K-L for representative images). PCR analysis of DG of male mice demonstrated that SK3 also impacts on the expression of specific genes related to adult neurogenesis. Male T/T mice showed an upregulation of Achaete-scute homolog 1 (ASCL1) (t(4)= 3.69, p<0.05), indicating that SK3 is possibly involved in the phenotype of type 2a cells, and the Neurotrophin-3 (NT3) (t(4)= 3.7, p<0.05), probably produced by mature granular neurons. The statistical analysis did not detect any significant changes in the mRNA expression of the genes SOX2, DCX, Microtubule-associated protein 2 (MAP2), Paired box protein-6 (PAX6), or Oligodendrocyte transcription factor-2 (OLIG2) (Figure 3M).

### 2.3 SK3 channels overexpression in male mice promotes an increased number of filopodia, dendritic spines and induces changes in NRG1/ErbB4 pathway in DG

Facilitation of adult hippocampal neurogenesis increased mushroom dendritic spines, and intracellular cascades involving neuroprotection are associated with a better cognitive performance. KCa channels influence the formation of dendritic spines (Ngo-Anh et al. 2005). In the dentate gyrus, SK3 channels are expressed in the dendritic spines of granule cell commissural/associational inputs and the entorhinal inputs (Ballesteros-Merino et al., 2014). Because our results indicated that SK3 overexpression preferentially induces changes in adult neurogenesis and intracellular pathways involved in neuroprotection in male mice, we have further evaluated the expression of other variables also involved in neuroplasticity.

Also, we could observe an increased number of immature filopodia spines in male SK3 transgenic mice, suggesting that SK3 channels might be involved in the maturation process of dendritic spines in granular neurons (t(7)= 2.44, p<0.05; Figure 5, A-F). Although Saito and colleagues (1992) have suggested that filopodia spines are an active part of the formation of synapses during neurodevelopment, in the adult brain this type of spine can be very chaotic leading to the formation of excessive and unstable synaptic connections, contributing to neuropsychiatric disorders (Jontes and Smith, 2000), such as Autism Spectrum Disorders (Durand et al., 2012). Notably, in Autistic Spectrum disorders, a dimorphic sex prevalence is found (approximately 4:1 male prevalence) (Zwaigenbaum et al., 2012). Besides, female autistic patients seem to have better memory performance than male patients (Zwaigenbaum et al., 2012).

Finally, given the important role of Neuregulin-1 (NRG-1)/Receptor tyrosine-protein kinase erbB-4 (ErbB4) in radial glia formation and neuronal migration, axon pathfinding and dendritic development (Lopez-Bendito et al., 2006, Rieff and Corfas, 2006), we investigated whether the genetic up-regulation of SK3 channels would change the pattern of expression of NRG-1 and the receptor tyrosine-protein kinase erbB-4 (ErbB4). Quantitative PCR analysis exhibited downregulation of NRG-1 mRNA in T/T male mice (Figure 5). No changes in ErbB4 expression (analyzed by WB and qPCR) were detected. The NRG1/ErbB4 signaling recruits PI3K/Akt, MAPK, Phospholipase C, and other pathways (Galvez-Contreras et al., 2013). The MAPK/ERK pathway has a central role in proliferation and survival cell regulation. The Epidermal Growth Factor Receptor is also regulated by Ca2 +-signaling, which could explain the adverse effects of SK3 overexpression on this pathway. Polymorphisms in NRG1 are associated with schizophrenia, autism, and may be involved in the pathophysiology of mood disorders and AD (Harrison and Law., 2006; Tian et al., 2017). SK3-overexpression-induced downregulation in NRG-1 gene expression would lead to defective ErbB4 receptor activation and, in consequence, to less recruitment of the intracellular pathways PI3K/Akt/mTOR and MAPK. Of note, both pathways are involved in the control of cognition, behavior, and in the facilitation of hippocampal neuroplasticity (Franco et al., 2017), events disrupted in SK3-T/T male mice.

### 2.4 SK3 overexpression induces sex-dependent downregulation of estrogen signaling, ERK1, JNK, and Akt/mTOR pathways in the DG of males but not in female mice

This sex dichotomy may rely on the changes in the estrogen signaling observed in the DG of male T/T mice. Several articles have shown significant increases in the proliferation and survival of newborn neurons in the hippocampus after treatment with estradiol (Barker and Galea, 2008; McClure et al., 2013). In ovariectomized rats, the proliferation of new neurons is significantly reduced, while estrogen replacement reverses the effect of ovariectomy (Pawluski & Galea, 2006; Pawluski et al., 2006).

In the hippocampus, estrogen might facilitate the induction of LTP. Studies also offer evidence that estrogen treatment in mid-age would improve aging-induced cognitive decline (Black et al., 2016). Based on the general hypothesis that estrogen actions in the hippocampus account for the sex dimorphism observed during memory process (McEween, 2002) and that levels of 17-β estradiol are related to SK3 expression levels, we sought the analysis of the levels of these sex hormones in the DG or hippocampus of WT or T/T mice (both sexes). Our results showed that as expected, female mice express higher levels of 17-β estradiol in DG and hippocampus than male mice, independently of the genotype. Likewise, this sex hormone levels did not differ between female WT and T/T mice in DG or the hippocampus – Figure 4B. On the other hand, male T/T mice expressed higher levels of 17-β estradiol than their WT controls in the DG (t(6)= 2.1, p=0.07) and hippocampus (t(6)= 8.4, p< 0.001)- Figure 4A.

**Figure 4.**
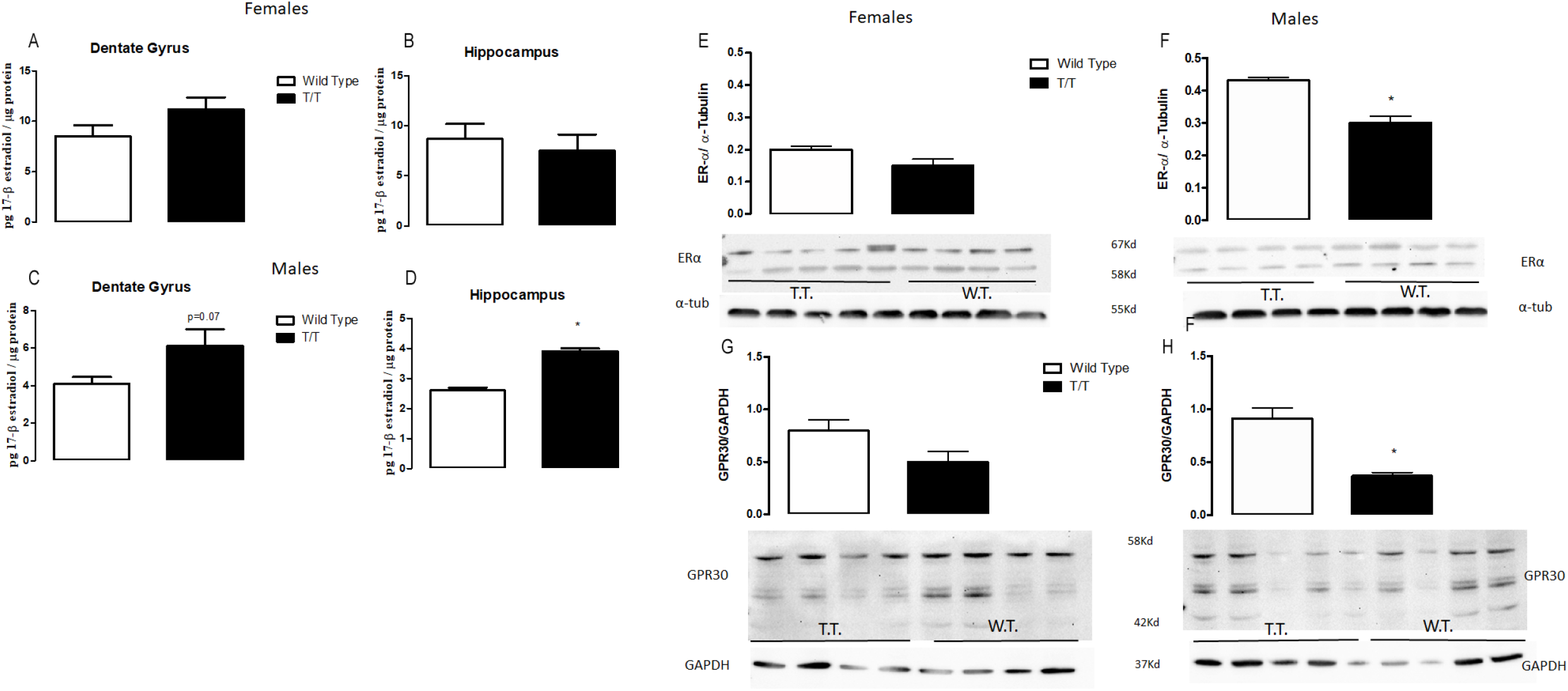
The estrogen signaling is dysregulated in the DG of T/T male, but not female, mice. A and C: levels of 17-β-estradiol in the dentate gyrus of female and male mice, respectively. B and D: levels of 17-β-estradiol in the hippocampus of female and male mice, respectively. E and G: expression of estrogen receptors ER-α and GPR30 in the DG of female mice determined by western-blot analysis. F and H: same as E and G but in male mice. Bars represent mean+/− SEM. N=5 per group.

**Figure 5.**
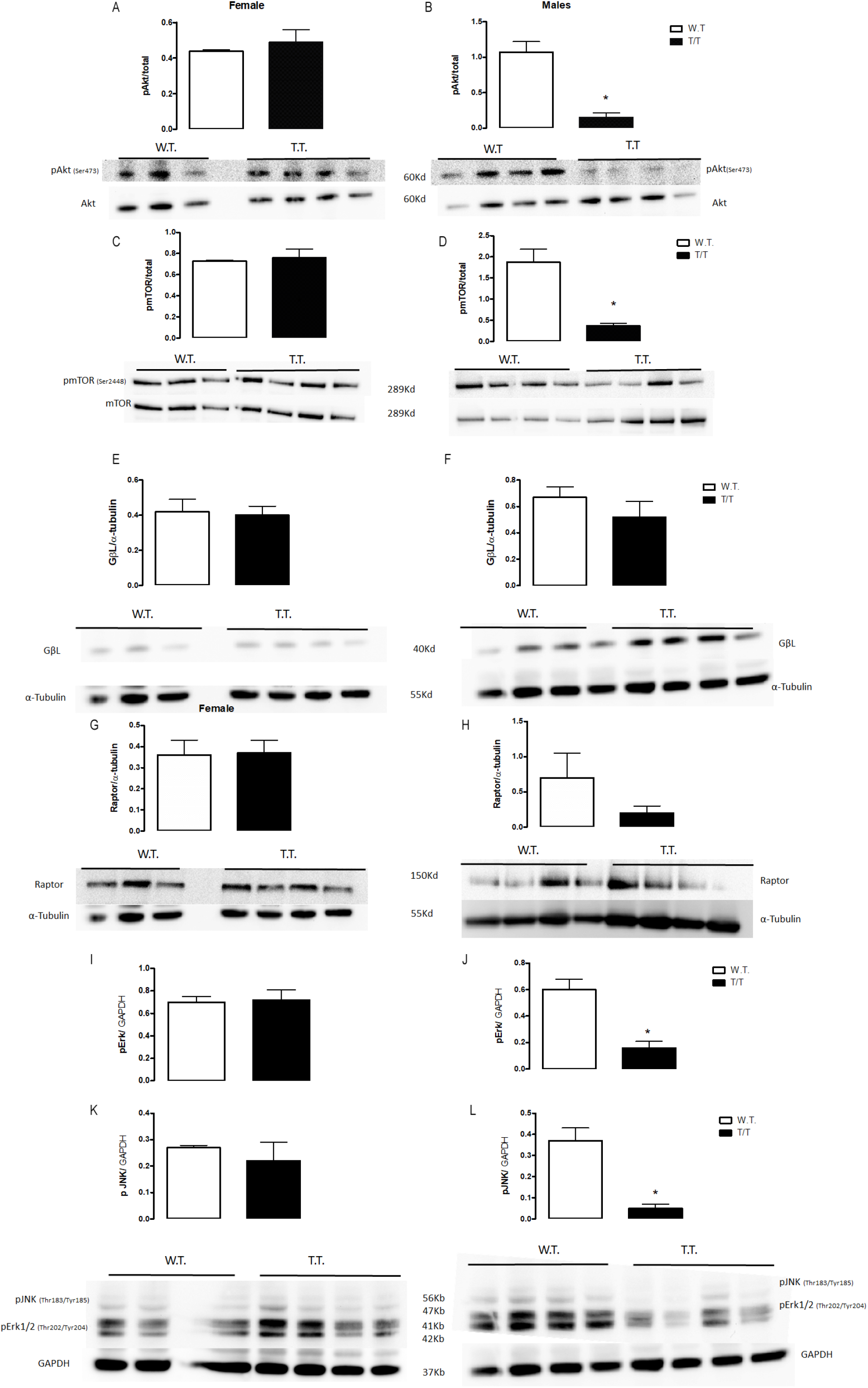
SK3 overexpression negatively modulates Akt/mTOR and MAPKinase, signaling the DG of male, but not female mice. A and B: pAkt/total Akt expression measures by Western blot (WB) in the DG of female and male mice, respectively. C and D: pmTOR/mTOR; E and F: GβL/α-tubulin; G-H: RAPTOR/α-tubulin; I-J: pERK1/2/GAPDH; pJNK/GAPDH. Representative images of the entire row of the WB membrane and its respective loading control. Bars represent mean+/− SEM. N=4 per group., except for the ERK and JNK in the WT group (n=3). * represents a difference from the WT group; p<0.001.

Estrogen levels highly modulate SK3 channel expression. In the brain, estrogen exposure induces SK3 mRNA expression in several brain regions (Bosch et al., 2002). Estrogen not only regulates SK3 channel expression but also can modulate its function. For instance, in GABAergic pre-optic neurons, estrogen enhanced the ability of the α-1 receptor to inhibit SK channels, increasing cell excitability (Kelly et al., 2002). Although not studied yet in the hippocampus, this mechanism (estrogen-induced SK3 inhibition) might counterbalance the mechanism by which KCa channels induced longer-lasting sAHP delaying a new event of neuron depolarization.

Curiously, male T/T mice present a lower expression of estrogen receptors in the DG of the hippocampus (ERalpha; t(6)= 5.2, p< 0.01; and GPR30 t(6)= 4.9, p< 0.01) than the WT (Figure 4C) despite their higher levels of estrogen, a possible compensatory mechanism. On the other hand, female T/T and their controls WT present similar expression profile of the aforementioned receptors (ER-a- t(7)= 0.8, p= 0.4; GPR30- t(7)= 1.2, p= 0.3)- Figure 4D. Interestingly, in male autistic patients, estrogen signaling seems to be disrupted (Crider et al., 2014). Like in T/T male mice, the expression of GPR30 is found to decrease in patients with Autistic Spectrum disorders (Altun et al., 2017).

Estrogen activates multiple signaling pathways and cellular processes via both genomic and non-genomic actions (Spencer et al., 2008). Recently, it was suggested that G-protein coupled estrogen receptors might improve cognitive outcomes in traumatic brain-injured rats via the PI3K/Akt pathway (Wang et al., 2017). Estrogens can also stimulate signaling and activation of the Erk-1/2-MAPKinase pathway, via Estrogen Receptor (ER)-α, ER-β, or GPR30 (G-protein coupled estrogen receptor 30 0r GPR1) (Fillardo et al., 2000). MAPK-pathway seems to be required for LTP formation during learning tasks (Giovannini et al., 2001).

Our western blot analysis suggested that changes in Akt/mTOR and MAPkinase pathways induced by SK3 overexpression rely on sex dimorphism (Figure 2). In agreement with the results of NOR, WB analysis of DG suggested reduced expression of phosphorylated forms of the extracellular signal-regulated kinase ½ (pERK) in male, but not female mice, (t(6)= 4.75, p<0.01; figure 2, A- male and B- female), Jun N-terminal kinase (pJNK) (t(6)= 5.11, p<0.01, figure 2 C and D); pAKT (t(6)= 5.32, p<0.01; figure 2 E and F), pmTOR (t(6)= 4.66, p<0.01, figure 2, G and H). Regarding Akt/mTOR pathway, our results are inconclusive about the role of SK3 on mTORC1 and mTORC2 pathways since no significant effects on the expression of RAPTOR and RICTOR were detected (data not shown). We hypothesize that the down-regulation of both PI3K/Akt/mTOR and MAPK/ERK1/2, MAPK/JNK is observed in male but not in female T/T mice, because estrogen signaling, would act as a compensatory and alternative pathway via estrogen receptors ER-a and GRP30. This estrogen-mediated event would allow female T/T mice to counterbalance the activation of MAPK and Akt/mTOR signaling even in the presence of SK3 overexpression.

In summary, our results suggest that SK3 participates in distinct forms of neuroplasticity in the dentate gyrus. We also found a connection between SK3 channels and estrogen signaling as a possible mechanism mediating the sex dimorphism of memory processes within the DG of the hippocampus.

The results presented here also generate data to support the importance of studying the effects of genetic manipulations in mice of both sexes. Unveiling this SK3-mediated mechanism on sex dimorphism help understand differences in the prevalence and in the course of symptoms of mental disorders that have already described sexual dimorphisms, such as autism, depression, and Schizophrenia (Koss & Frick, 2017).

## 3.0 Materials and methods

### Animals

Adult male and female (2,5–3 months old) wild type (WT) and homozygous SK3 (SK3-T/T) littermate mice were used for all analyses (Martin et al., 2016; Grube et al., 2011). The SK3 conditionally overexpressing mice (Bond et al., 2000) were kindly provided by John Adelman and Chris Bond. In our manuscript, we have used the “T” in reference to the overexpressing allele of SK3 (homozygotes are indicated as “T/T”). All animals were housed in group and kept in a temperature-controlled room with free access to water and food during all experimental procedures. We used only female mice without regard to their estrous cycle since we were unable to accurately discern the various stages by visual observation. After weaning, all littermates were separated by sex and kept in microisolators and separate rooms during all the procedures. Also, experimental females were never exposed to soiled bedding from male cages, which could synchronize their estrous cycle (Campbell et al., 1976; Caligione, 2009). Besides, exposure to male pheromones might influence females’ performance in memory tests and enhance adult neurogenesis (Mak et al., 2007). For the experiments involving BrdU (Sigma, St. Louis, Missouri-dissolved in 0.9 % NaCl and filtered sterile at 0.2 m), mice received two doses of 100 μg/g body weight at a concentration of 10 mg/ml(intraperitoneally) -one injection per day, for two consecutive days. All experiments were approved by the local Animal Ethics Committee (No. AZ 33.9-42502-04-10/0314) and performed according to German laws.

### Behavioral procedures

#### Novel object recognition test

In the new object recognition test (NOR), animals were allowed to explore a round arena (open field) in 3 different moments. The task did not involve any physical suffering except for the mild stress promoted by the exposure to a novel environment and human manipulation. On the first day, animals explored the arena for 5 minutes (habituation session). 24 hours later, a training session was conducted. During this session, animals were again placed in the open field by the experimenter, and they were free to explore the arena again for another 10 minutes. However, this time they were exposed to two identical objects (object A1 and A2; Lego Toys) positioned in two adjacent corners (distanced 8 cm from the walls). The short-term retention memory (STM) was conducted 1.5 hours after the training session, whereas the long-term retention memory (LTM) test took place 24 hours after the training session. During the test session, mice explored the open field for 5 minutes in the presence of the familiar (A) and a novel (B or C) object. Objects (new and old) are distinct in shape and color (although made of the same material-plastic). The exploratory preference was defined according to Bevins and Besheer (2006) as the percentage of the total time that animals spent investigating the new object. It was calculated by the ratio TN / (TF+TN) * 100 (where TN = time spent exploring the novel object; TF= time spent exploring the familiar object).

#### SocioBox test

To analyze the influence of SK3 overexpression on social recognition memory, animals were tested in the SocioBox (Krueger-Burg et al., 2016). This paradigm tests the social recognition abilities of mice exposed to a familiar cage mate among stranger ones. The SocioBox apparatus is composed of a circular behavioral chamber (Max Planck Institute of Experimental Medicine), made up of a grey plastic ground plate (diameter 56 cm) and an outer ring with five rectangular chambers, called inserts (width 8.5 cm, length 11.5 cm, height 20.5 cm) separated by grey plastic walls. The inserts are replaceable and have three grey plastic walls and a transparent removable plastic door which faces the open area in the inside. The transparent door consists of 31 holes (diameter 0.8 cm), which allow sensorial (odor) interaction between mice tested. Next to the apparatus, a white opaque plastic cylinder (diameter 19 cm, height 20.5 cm) was provided to separate the experimental mouse in the open area visually and spatially from the stimulus mice in the inserts during the initiation stage. As ‘stimulus mice’, which interact with the experimental mice, the C3H strain wild type animals were used because they have been reported to show robust social interaction during test situations (Moy et al., 2007).

Above the SocioBox, an overhead camera (Panasonic HD-Camcorder HC-V160) was fixed. The camera was turned on before lifting the cylinder and recorded for 5 min with a resolution of 25 frames per second. The videos were analyzed using the automated software Viewer3 (Biobserve). The software tracks the coordination points of the head, body, and tail of the mouse, as well as the time the experimental mouse spent in front of the inserts with the stimulus mice.

After three days of habituation, the test session took place. In the beginning, the experimental mice and stimulus mice were exposed separately to the desired light conditions for at least 30 min. The test session was divided into three phases – exposure 1, exposure 2, and memory test. The 10 min phases are composed of an initiation stage with the opaque cylinder in the open area (5 min) and an interaction stage without the cylinder (5 min). Six selected stimulus mice were placed in their inserts, and five of them were located in the SocioBox apparatus. All mice stayed in the inserts for the same time, since the activity of the mice decreased over time. The initiation stage in which the experimental mouse was placed in the cylinder inside the SocioBox apparatus for 5 min took place next. With the overhead camera recording, the interaction stage of exposure 1 started when the cylinder was lifted to allow interaction with the stimulus mice. After 5 min, the experimental mouse was placed in the transportation cage while cleaning the SocioBox apparatus and changing the transparent doors as described in the habituation session. In the next step, the experimental mouse was put back in the cylinder for exposure 2. Finishing the second cleaning procedure, one stimulus mouse (Sori) was removed from the SocioBox apparatus, and the stranger mouse (S new) was placed in the slot. The position of the stranger mouse was varied to avoid spatial bias. In the end, all mice were put back in their home cages, and the SocioBox apparatus was cleaned with 70 % ethanol to prepare it for the next test session.

### Experimental Design

#### Experiment 1

Animals with mixed genotypes (male and female) were housed in groups of 3-5 per cage. A total of 20 animals were used (10 animals per group of each genotype- WT or T/T). Animals were submitted to the NOR test and, one week later, to the PPI analysis (described in the Supplementary material). Of note, male and female mice were tested in different days/experiments to avoid interference of smell and sexual behavior in the tests.

#### Experiment 2

Experiments were performed with animals of C57BL/6J background as ‘experimental mice’. At the beginning of the testing, the mice were 14 to 16 weeks old. C3H WT animals of the same gender and age were used as ‘stimulus mice’. All experiments were conducted during the light phase of the day (8 am to 6 pm) with a light intensity of 10– 15 lux. The investigator was blinded to the genotype of the experimental mice.

#### Experiment 3

Similar to the experiment 1, but animals were submitted to tail suspension and sucrose preference test (described in the supplementary material). Animals (n=10/group) received two injections of BrdU (100 mg/Kg) and were left undisturbed in their home cages for 11 days. On 11^th^ day after BrdU injection, animals were submitted to the tail suspension test and one week later to the sucrose preference test.

Animals were then divided into two groups: Group 1- One day after the sucrose preference test, the first group were decapitated under deep anesthesia (Ketamine-60mg/Kg and, Xylazine 10mg/Kg), the brains removed and the dentate gyrus dissected for Western Blotting (WB). Group 2-animals were perfused 20 days after the last BrdU injection under deep anesthesia and their brains prepared for immunofluorescence protocols (BrdU-neurogenesis analysis).

### Immunohistochemistry experiments

#### Tissue preparation

Twenty days after the last BrdU injections, mice were anesthetized with ketamine/xylazine (60 mg/kg – 8 mg/Kg) and perfused transcardially with 25 ml of phosphate buffer saline (PBS) 0.1 M (pH=7.4) followed by 25 ml of 4 % paraformaldehyde. The brains were removed, immersed in the same fixative solution for 12 h, and then transferred into a 30 % sucrose solution (72 h), and frozen in isopentane (−30 °C). Cryostat-cut coronal sections (30 μm) were collected (rostral-caudal sections of the whole hippocampus) and then submitted to immunofluorescence/histological procedures.

#### Immunohistochemistry for Doublecortin (DCX)

Coronal free-floating brain sections (30 μm) were processed as described elsewhere (Campos et al., 2013). Briefly, after 1 hour of blockade (PBS + 0.25 % Triton X-100 + 1 % bovine serum albumin -BSA), brain sections were incubated overnight at 4 °C with primary antibodies (goat polyclonal anti-doublecortin- Santa Cruz Biotechnology, California, USA) followed by incubation for 1 h at room temperature with the appropriate highly cross-adsorbed secondary antibodies (Invitrogen, California, USA). Immunoreactivity was detected by the avidin-biotin immunoperoxidase method (Vectastain ABC kit, Vector Lab, Burlingame, CA, USA), and the product of the reaction was revealed by adding the chromogen 3,3-diaminobenzidine tetrahydrochloride (Sigma Chemical, St. Louis, USA). Doublecortin migrating cells were quantified in a minimum of 6 coronal sections per animal. Positive cells were quantified in the granular layers of the dentate gyrus of the hippocampus (400x magnification) and the number of positive cells estimated by calculating the total hippocampal volume as determined by the sum of the areas of the sampled sections multiplied by the distances between them (series of hippocampal sections located between 1.3 mm and 2.5 mm posterior to bregma – Paxinos and Franklin, 2006). Doublecortin migrating cells were quantified using a computerized image analysis system (ImagePro software) coupled to an Olympus light microscopy (Bx60) (Campos et al., 2013).

#### Immunofluorescence for BrdU detection

Coronal brain sections (30 μm) were transferred to a 24-well plate (free-floating at 8/well) and then washed three times with PBS. For BrdU analysis, slices were incubated for 30 min with HCl (2N) at 37 °C and cooled for 10 min at 4 °C. After, slices were washed 3 times in Boric Acid solution (pH= 8.9) and, subsequently, 3 times in PBS. After these procedures, slices were blocked for 2 hours in 1 % of Bovine Serum Albumin (BSA) + 0.25 % Triton X-100 in PBS solution. After the blocking period, slices were incubated overnight at 4 °C with the respective primary antibodies (rat-monoclonal anti- BrdU −1:200- Abcam; Rabbit polyclonal antibody anti-DCX,1:500- Cell Signaling; mouse monoclonal anti-NeuN- Millipore) diluted in a 0.1 % BSA and 0.125 % Triton X-100 solution. Subsequently, they were washed 3 times in PBS and then incubated with the specific secondary antibodies (Alexa fluor 488, 594 and 647- 1:1,000; Invitrogen) for 2 hours at room temperature. Slices were mounted onto glass slides for microscopic analysis. The staining (BrdU; BrdU/DCX; BrdU/NeuN and BrdU/DCX/NeuN were analyzed using a confocal microscope Leica TSC-SPE (Leica) with two passes by Kalman filter and a 1024 × 1024 collection box. For BrdU- and doublecortin-positive cells were quantified in the subgranular zone of the hippocampus, and a minimum of 6 slices were analyzed (2 dentate gyrus/animal-distributed rostrocaudal between 1.3 and 2.5 mm posterior to bregma; Paxinos and Franklin, 2006). For double and triple staining, we followed 100 cells using z-stacks (10 mm scan; 10 steps) projected in 3D by Leica software to determine co-labeling (400x magnification). Positive cells were normalized to the dentate gyrus area determined with a 10x objective, and the absolute number of positive cells was calculated considering the total hippocampal volume as determined by the sum of the areas of the sampled sections multiplied by the distances between them (Campos et al., 2013).

#### Western Blotting

Animals were sacrificed, and the DG dissected as previously described elsewhere (Walker and Kempermann, 2014). DGs were homogenized in a lysis buffer (NaCl 137 mM, Tris-HCl 20 mM pH= 7.6, 10% glycerol, NaF 100 mM, NaVO3 10 mM, 0.1 % Triton-X-100) and protease inhibitor cocktail (Roche- Basel- Switzerland) for protein extraction. After two centrifugation steps (12000 rpm, 10 min, 4°C), the supernatants were preserved and stored at −80°C. The total protein of the samples was determinate by BCA protein assay (Roti®-Quant universal – Carl Roth). Homogenates were submitted to an SDS-PAGE protein separation in 4-12 % or 3-8 % (depending on the molecular weight of the protein of interest) NuPAGE bis-tris gels (Invitrogen) and transferred to a nitrocellulose membrane (Amersham). After the transference, membranes were washed in a solution of TBS+0.01 % Tween-20 (TBS-T) and incubated in 5 % non-fat milk (Biorad) blocking solution for 2 h. Soon after, they were incubated overnight (4 °C) with the primary antibodies (Cell Signaling, Danvers-MA-USA: rabbit anti-phospho-Akt Ser473 1:1000, rabbit anti-Akt 1:1000; rabbit anti-phospho-mTOR2448 1:1000, rabbit anti-phospho-mTOR2481, mouse anti-mTOR 1:1000, rabbit anti-Raptor 1:1000, rabbit anti-Rictor 1:1000, rabbit anti-GβL 1:1000, rabbit anti-phospho-ERK1/2 1: 1000, rabbit anti-phospho JNK 1:1000; Millipore, Darmstadt, Germany, rabbit anti-Estrogen Receptor α; 1:500; Abcam, Cambridge-United Kingdom, Anti-G-protein coupled receptor 30 1:500). After overnight incubation, membranes were washed 3 times in TBS-T and incubated with the specific secondary antibodies for 1 h (anti-mouse or anti-rabbit HRP 1:1000, Amersham, Buckinghamshire, United Kingdom) and revealed by a chemiluminescent method (ECL, Millipore). Unsaturated, normalized and preserved bands had their density quantified using the Software LICOR Image Studio Digits (version 3.1, Lincoln, Nebraska USA)

#### 17-β-estradiol levels determination by ELISA

The levels 17 β-estradiol were quantified in DG and CA1/CA3 homogenates derived from an independent group of animals (male and female, T/T, and WT mice) were detected by ELISA colorimetric detection kit (Enzo Biochem- Farmingdale, New York-USA) according to the manufacturers’ instructions. Results obtained were controlled by total protein concentrations determined by BCA assay.

#### Golgi staining analysis

An independent group of male animals (not submitted to behavioral tests) was sacrificed by cervical displacement under deep anesthesia (ketamine 60 mg/Kg and xylazine 8 mg/Kg) and their brains removed. The Golgi-Cox procedure was performed using the FD Rapid Golgi Stain Kit (FD Neurotechnologies- Columbia, USA) accordingly to the fabricant’s recommendations. At the end of the process, brains were rapidly frozen in isopentane and dry ice and cut at 100-μm thick sections in a cryostat (Cryocut 1800, Leica, Heerbrugg- Switzerland) to obtain slices containing the hippocampus. The sections were transferred to gelatinized slides, and two days after drying at room temperature, they were stained and submitted to alcohol-induced dehydration. Finally, the slides were covered with Permount™ (Fisher Scientific-Hampton, USA) and analyzed in confocal microscopy following the method described elsewhere (Spiga et al., 2011). An experimenter blinded to the groups performed the analysis. Golgi-Cox impregnated neurons located into the dentate gyrus were localized using the transmitted light function of the confocal microscopy (TSC-SPE- 20x objective). Afterward, they were scanned in the confocal microscopy by setting up the reflection mode, using the 488 nm wavelength and replacing the dichroic filter by a 30/70 beam splitter (Spiga et al., 2011). The number of dendritic spines in 10 μm-branches classified as secondary and tertiary were counted in six neurons per animal using the following criteria: the neurons were relatively isolated, displayed a defined cell body and presence of intact dendritic three and primary, secondary and tertiary branches. Spines were then classified as thin, stubby, filopodia, or mushroom, according to their morphological features.

### Laser capture microdissection and *quantitative real-time (qRT)-PCR* analysis

#### Preparation of brain cryo-sections for laser capture microdissection

animals were sacrificed by CO2 inhalation and subsequently decapitated. Brains were frozen on dry ice and afterward kept at −80 °C. Cryosections of male WT and SK3-T/T mice (n=3) were mounted on polyethylene naphthalate (PEN) membrane slides. For dehydration, sections were subsequently fixed two times in 70% EtOH for 30 sec, followed by a dehydration series in 95% EtOH for 30 sec, two times’ freshly’ purred with 100% EtOH for one minute and two times xylene for two minutes. After dehydration, slides were air-dried, transported, and kept in a box with desiccant. We used only precooled RNAse free plastic ware and solutions.

#### Laser capture microdissection of the dentate gyrus and qRT-PCR

The dentate gyrus of the dorsal hippocampus region (Bregma −1.34 mm until Bregma −2.46 mm) were microdissected from 10–15 adjacent slides (20 μm) of each animal and were pooled. For the microdissection, we used the Arcturus Veritas microdissection system. For cutting a UV laser (Power = 5–10%) and for attachment, an IR laser (Power: 80−90 mW and pulse duration: 2500–5000 μs) were used. Successful cutting and collection steps were subsequently validated in bright-field on the quality control slot of the device. Microdissected samples were lysed in 100 μl of RNA lysis buffer (Qiagen, Hilden, Germany) by vigorous vortexing for 30 sec and stored at −80°C until further use. All procedures were done under RNAse-free conditions. For isolation of minimal amounts of total RNA (100 pg—100 ng), the RNeasy micro kit (Qiagen) was used according to the manufacturer’s instructions. Quality and concentration of RNA were examined with the Nano LabChip and the Bioanalyzer2000 (Agilent Technology, Santa Clara, USA). The first-strand cDNA from total RNA was synthesized using oligo-dT as a primer (SuperScript; Life Technologies, Darmstadt, Germany) according to the manufacturer’s instructions. The cDNA was stored at −80°C for further use. The qRT-PCR was performed in triplicate using the TaqMan system in the LightCycler 480 (Roche, Mannheim, Germany).

The relative abundance of SOX2, GFAP, ASCL1, DCX, MAP2, PAX6, OLIG2, NTF3, NRG1, and ERBB4 transcripts in the dentate gyrus of the mouse brain was studied by quantitative RT-PCR. The Ct value of these target genes was normalized to the reference genes hypoxanthine-guanine phosphoribosyltransferase 1 (HPRT1).

For quantifying mRNA expression by real-time PCR, the following fragments were amplified: nt 134–233 from sequence NM_013556 detected with the mHPRT1 probe (5′-(Fam)-CAGCGTCGTGATTAGCGATGATGAACCAGG-(Tamra)-3′); nt 172–244 from sequence NM_011443 detected with the mSOX2 probe (Universal ProbeLibrary probe #63, Roche, Cat.No. 04688627001; 5′-(FAM)-AGGAGGAG-(Dark Quencher)-3′); nt 806-905 from sequence NM_010277 detected with the mGFAP probe (5′-(Fam)-TGGTATCGGTCTAAGTTTGCAGACCTCAC-(Tamra)-3′); nt 1280-1350 from sequence NM_008553 detected with the mASCL1 probe (Universal Probe Library probe #74, Roche, Cat.No. 04688970001; 5′-(Fam)-GGCAGCAG-(Dark Quencher)-3′); nt 588-682 from sequence NM_001110222 detected with the mDCX probe (Universal ProbeLibrary probe #68, Roche, Cat.No. 04688678001; 5′-(Fam)-CTGCTCCT-(Dark Quencher)-3′); nt 118-230 from sequence NM_001039934 detected with the mMAP2 probe (5′-(Fam)-TGCTGTGTGCTCCAAGTTTCA-(Tamra)-3′); nt 225-323 from sequence NM_001244198 detected with the mPAX6 probe (Universal ProbeLibrary probe #67, Roche, Cat.No. 04688660001; 5′-(Fam)-CTCCAGCA-(Dark Quencher)-3′); nt 33-140 from sequence NM_016967 detected with the mOLIG2 probe (Universal ProbeLibrary probe #21, Roche, Cat.No. 04686942001; 5′-(Fam)-CAGAGCCA-(Dark Quencher)-3′); nt 376-486 from sequence NM_008742 detected with the mNT3 probe (Universal ProbeLibrary probe #49, Roche, Cat.No. 04688104001; 5′-(Fam)-GGCCACCA-(Dark Quencher)-3′); nt 757-864 from sequence NM_178591 detected with the mNRG1 probe (5′-(Fam)-TGTGCAAGTGCCCAAATGAGTTTACTGGT-(Tamra)-3′); nt 828-938 from sequence NM_010154 detected with the mERBB4 probe (5′-(Fam)-GATGTGGAAGATGCCATAAGTCTTGCAC-(Tamra)-3′). Conditions for PCR were 2 min at 50 °C, 10 min at 95 °C, 15 s at 95 °C, 15 s at 56 °C, and 1 min at 60 °C (50 cycles). Calculations of gene expression were performed using the ∆∆Ct method, according to Pfaffl and colleagues (2001).

### Statistical analysis

Data obtained were tested for normality (Kolmogorov-Smirnov’s Test), homogeneity of variance (Levene’s test), and then analyzed by Student’s t-test. The significance level was set at p < 0.05. Data were expressed as mean ± SEM. Data from the SocioBox experiments were analyzed by One-Way ANOVA. Statistical analysis was performed by SPSS version 16.0 software (IBM, USA).

## 5.0 Acknowledgments

Thank you, Sabine Stolpe, Tanja Nilsson, Ursula Kutzke, Sabine Klöppner, Araceli Sabchez, Eleni Tamburus, and Flavia Turcato, for their exceptional technical support. We give special thanks to John P. Adelman and Chris T. Bond, Vollum Institute, OHSU Portland, OR, for providing the SK3-T/T mice. We also acknowledge the work of the staff of the animal facility at the Max Planck Institute for Experimental Medicine. Special thanks to Professor Andre Fischer for allowing us to perform the PPI test in his group’s behavioral facility at the European Neuroscience Institute Göttingen (ENI-G). The Max Planck Society and CAPES-DAAD financially supported this work. ACC was a recipient of CAPES-DAAD fellowship (#3089/14-2) and is currently supported by a FAPESP fellowship (2015/05551-0) and CNPq (type 2 research productivity fellowship). SM was supported by the Cluster of Excellence and DFG Research Center Nanoscale Microscopy and Molecular Physiology of the Brain. FFS has a FAPESP scholarship (2019/09178-3). We FAPESP, CNPQ, and CAPES-PG.

## 6.0 Authors contribution

ACC designed the experiments, performed all behavioral experiments (except the SocioBox), all WB and immunostaining analysis and, wrote the first draft of the manuscript. FFS performed and analyzed the immunofluorescence labeling and the Golgi staining technique. SM, ML have helped in behavioral test, WB and performed laser dissection, qPCR and immunostaining protocols. DH performed the SocioBox test. MASS performed the corticosterone analysis. LP, WS, and EDB were responsible for the grants that allowed carrying out all experiments presented here, as well as helped to design and discuss the experiments and their results. All authors have read and approved the final version of the manuscript.

## Supplementary Material and Methods

### Prepulse inhibition (PPI)

The test took place in soundproof chambers. The chambers contain an inner chamber equipped with a loudspeaker and startle platform (accelerometer) linked to a computer (TSE GmbH, Bad Homburg, Germany). Mice were placed individually in small metal grid cages (90 × 40 × 40 mm) to restrict major movements and exploratory behavior. The experimental session consisted of a 3 min habituation to 65 dB background white noise (continuous throughout the session), followed by a baseline recording for 2 min. For PPI tests, a non-startling prepulse stimulus (70, 80 or 90 dB) and frequency of 7,000 Hz (duration: 20 ms) were applied before the startle stimulus (white noise of 120 dB of 40 ms; in a pseudorandom order with inter-trial intervals ranging from 8 to 22 s). An interval of 100 ms with background white noise was set between each prepulse and pulse stimulus. The amplitude of the startle response (expressed in arbitrary units, AUs) is defined as the maximum force detected as a reaction to the acoustic stimulus detected by the sensors attached to the cage. PPI (expressed in %) is calculated as the percentage of the startle response using the following formula: PPI (%) = 100 − [(startle amplitude after prepulse and pulse)/(startle amplitude after pulse only) × 100] (Issy et al., 2009). Each experiment lasts around 25 min per mouse in which 76 different events will be presented: 7 events of habituation pre-experiment 65 dB background white noise; 8 events of white noise 120 dB; 12 events each of PPI 70 dB; PPI 80 dB or PPI 90 dB followed by a white noise presentation of 120 dB; 15 events of no stimulus and 10 events of habituation post-experiment (5 min) under a 65 dB background white noise.

### Tail suspension test (TST)

This test was conducted to screen passive and active coping behaviors that are modified negatively by stress and positively by antidepressant drugs. The TST is based on the premise that animals subjected to inescapable stress, promoted by the suspension of their tails, develop an immobile posture. The total duration of immobility was measured according to the method proposed by Steru et al. (1985) in a 6-minute session. Mice were suspended 50 cm above the floor by their tails (fixed with adhesive tape) and the immobility time scored during the total time of the test session (6 minutes). A video camera recorded the test, and an investigator blind to treatment/genotype of the animals counted the percentage of time spent immobile during a 6-minute session.

### Sucrose preference test

This reward-based test measures, in rodents, behaviors related to anhedonia, a symptom observed in patients diagnosed with a depressive disorder characterized by the loss of interest and inability to feel and enjoy pleasurable situations. For the sucrose preference test, animals were isolated 24 hours before the test and exposed to two bottles containing plan water in their new home cages. Twelve hours before the test, the animals were food-deprived. The sucrose preference test was carried out in the home cages of experimental animals. Briefly, we presented them with two bottles of the same size, color and material, one containing plain drinking water and a second one filled with a 2 % sucrose solution (Willner et al., 1987). The test duration was overnight. At the end of the test, the percentage of sucrose preference was calculated by the following formula: [amount of sucrose intake (mL)]/ [amount of sucrose intake (mL) + amount of water intake (mL)] * 100.

### Hemotoxylin-eosin staining and SK3 immunolabeling

Protocol used were very similar to the one described for the immunostaining of DCX using DAB-peroxidase method (SK3 antibody; 1:500- Alomone Labs- Jerusalem, Israel).

### SK3 expression by WB

similar to the protocol described for the determination of pAkt and total Akt (SK3 antibody; 1:500-Alomone Labs-Jerusalem, Israel)

### Corticosterone assay

Blood samples were taken after euthanasia under deeps anesthesia. Samples were centrifuged at 4 °C for 10 min with a speed of 4000 rpm and corticosterone plasma concentrations by ELISA (IBL International, Hamburg, Germany) (Houston et al., 2016).

### Laser capture microdissection of CA1 and qRT-PCR

the same protocol used for DG analysis. Additional genes were analyzed (Caspase 3- CASP3; SK3-KCNN3; S100, Dopamine receptor 2, CREB, BDNF, and mAPP).

## Legends for Supplementary figures

**Supplementary figure 1-** SK3-induced hippocampal shrinkage in male and female mice. A and B: Hematoxylin-Eosin stain (40x). Left panel T/T mice, right panel WT mice. C and D: expression of SK3 in the hippocampus and DG of T/T (left panel) and WT (right panel) mice (100x). F and G: WB expression of SK3 in the hippocampus. Red rectangles highlight the overexpressed SK3-protein chain. H and I: same as F and G but in the hippocampus. The bar represents 40uM.

***Supplementary figure 2-*** SK3 overexpression induces antidepressant-like behavior but does not change prepulse inhibition. A and B: prepulse inhibition in male and female mice, respectively. C and D: percentage (%) of immobility time measured during the tail suspension test. F and G: % of preference for sucrose over water. Bars represent mean+/− SEM (N= 10 per group); p< 0.05.

***Supplementary figure 3-*** Serum corticosterone levels and relative expression of genes related to neuroplasticity in the CA1 region of the hippocampus of WT and T/T male mice. Left upper panel: serum levels of corticosterone in female mice; Left lower panel serum levels of corticosterone in male mice. Right panel: the normalized ratio of mRNA expression of SK3, Olig2, CASPase-3, MAP-2, S100B, GFAP, NRG1, ErBB4, DRD2, BDNF, and mAPP (N=3/group). Bars represent mean+/− SEM. * indicates differences from WT. *p< 0.05 and **p< 0.01.

